# Genome-wide association analyses in > 119,000 individuals identifies thirteen morningness and two sleep duration loci

**DOI:** 10.1101/031369

**Authors:** Samuel E. Jones, Jessica Tyrrell, Andrew R. Wood, Robin N. Beaumont, Katherine S. Ruth, Marcus A. Tuke, Hanieh Yaghootkar, Youna Hu, Maris Teder-Laving, Caroline Hayward, Till Roenneberg, James F. Wilson, Fabiola Del Greco, Andrew A. Hicks, Chol Shin, Chang-Ho Yun, Seung Ku Lee, Andres Metspalu, Enda M. Byrne, Philip R. Gehrman, Henning Tiemeier, Karla V. Allebrandt, Rachel M. Freathy, Anna Murray, David A. Hinds, Timothy M. Frayling, Michael N. Weedon

**Author notes:** Joint contribution (Timothy M. Frayling, Michael N. Weedon). **Corresponding author:** Michael N Weedon, University of Exeter Medical School, RILD building Level 3, Royal Devon & Exeter Hospital, Barrack Road, Exeter, EX2 5DW.

## Abstract

Disrupted circadian rhythms and reduced sleep duration are associated with several human diseases, particularly obesity and type 2 diabetes, but little is known about the genetic factors influencing these heritable traits. We performed genome-wide association studies of self-reported chronotype (morning/evening person) and self-reported sleep duration in 128,266 White British individuals from the UK Biobank study. Sixteen variants were associated with chronotype (*P*<5x10^-8^), including variants near the known circadian rhythm genes *RGS16* (1.21 odds of morningness [95%CI 1.15, 1.27], *P*=3x10^-12^) and *PER2* (1.09 odds of morningness [95%CI 1.06, 1.12], *P*=4x10^-10^). The *PER2* signal has previously been associated with iris function. We sought replication using self-reported data from 89,823 23andMe participants; thirteen of the chronotype signals remained significant at *P*<5x10^-8^ on meta-analysis and eleven of these reached *P*<0.05 in the same direction in the 23andMe study. For sleep duration, we replicated one known signal in *PAX8* (2.6 [95%CIs 1.9, 3.2] minutes per allele *P*=5.7x10^-16^) and identified and replicated two novel associations at *VRK2* (2.0 [95% CI: 1.3, 2.7] minutes per allele, *P*=1.2x10^-9^; and 1.6 [95% CI: 1.1, 2.2] minutes per allele, *P*=7.6x10^-9^). Although we found genetic correlation between chronotype and BMI (rG=0.056, *P*=0.048); undersleeping and BMI (rG=0.147, *P*=1x10^-5^) and oversleeping and BMI (rG=0.097, *P*=0.039), Mendelian Randomisation analyses provided no consistent evidence of causal associations between BMI or type 2 diabetes and chronotype or sleep duration. Our study provides new insights into the biology of sleep and circadian rhythms in humans.

## Introduction

There are strong epidemiological associations among disrupted circadian rhythms, sleep duration and disease. A circadian rhythm refers to an underlying 24-hour physiological cycle that occurs in most living organisms. In humans, there are clear daily cyclical patterns in core body temperature, hormonal and most other biological systems ^1^. These cycles are important for many molecular and behavioural processes. In particular, circadian rhythms are important in regulating sleeping patterns. While each individual has an endogenous circadian rhythm, the timing of these rhythms varies across individuals. Those with later circadian rhythms tend to sleep best with a late bedtime and late rising time and are often referred to as an “owl” or as an “evening” person. Those with earlier rhythms tend to feel sleepy earlier in the night and wake up early in the morning and are referred to as a “lark” or “morning” person. The remainder of the population falls in between these extremes. This dimension of circadian timing, or chronotype, is one behavioural consequence of these underlying cycles. Chronotype can be simply assessed by questionnaire and is considered a useful tool for studying circadian rhythms ^2, 3^.

There is substantial evidence for a relationship between short sleep duration, poor quality sleep and obesity and type 2 diabetes ^4, 5^. Eveningness has been associated with poor glycaemic control in patients with type 2 diabetes independently of sleep disturbance ^6^ and with metabolic disorders and body composition in middle-aged adults ^7^. There is evidence from animal models that disruption to circadian rhythms and sleep patterns can cause various metabolic disorders ^8-10^. For example, mice homozygous for dominant negative mutations in the essential circadian gene, *Clock*, develop obesity and hyperglycaemia ^10^ and conditional ablation of the *Bmal1* and *Clock* genes in pancreatic islets causes diabetes mellitus due to defective β-cell function ^9^. Despite this evidence, in humans the causal nature of the epidemiological associations between sleep patterns, circadian rhythms and obesity and type 2 diabetes is unknown. Identifying genetic variants associated with sleep duration and chronotype will provide instruments to help test the causality of epidemiological associations ^11^.

A previous genome-wide association study (GWAS) in 4251 individuals identified a single genetic variant in *ABCC9* associated with sleep duration ^12^. A subsequent GWAS meta-analysis including 47,180 individuals identified a single locus for sleep duration near *PAX8* ^13^. There have been no published reports of variants influencing chronotype. The UK Biobank is a study of 500,000 individuals from the UK aged between 37 and 73 years with genome-wide SNP analysis and detailed phenotypic information, including chronotype and sleep duration (http://www.ukbiobank.ac.uk/). The UK Biobank study provides an excellent opportunity to identify novel genetic variants influencing chronotype and sleep duration which will provide insights into the biology of circadian rhythms and sleep and help test causal relationships between circadian rhythm and metabolic traits including obesity.

## Results

### Sixteen loci associated with chronotype in UK Biobank

Using self-reported “morningness”, we generated a binary and a continuous chronotype score. We performed genome-wide association studies on 16,760,980 imputed autosomal variants. **Figure 1** presents the overall results for these GWAS. **Table 1** presents details of all 16 genome-wide significant chronotype-associated loci.

**Figure 1.**
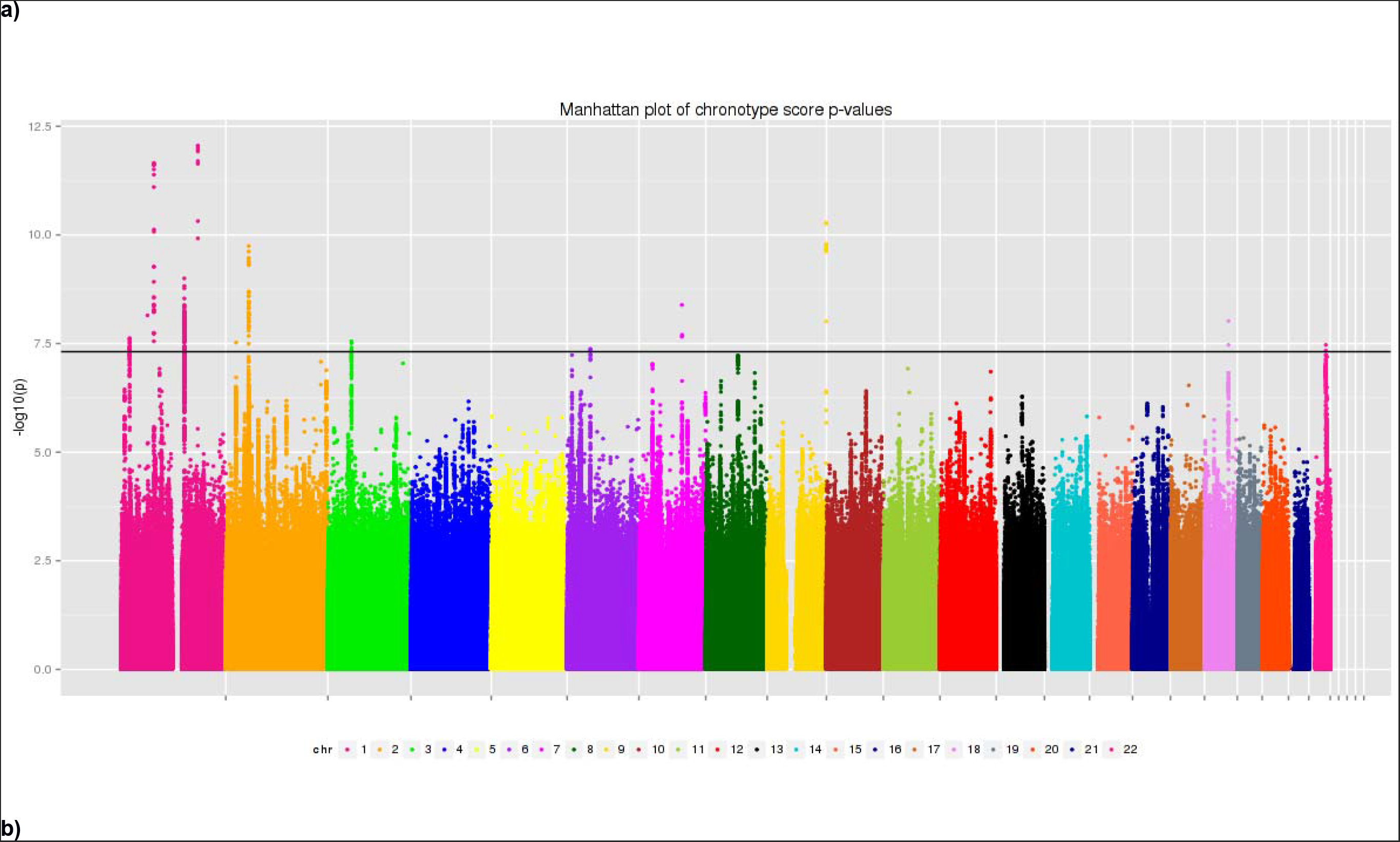
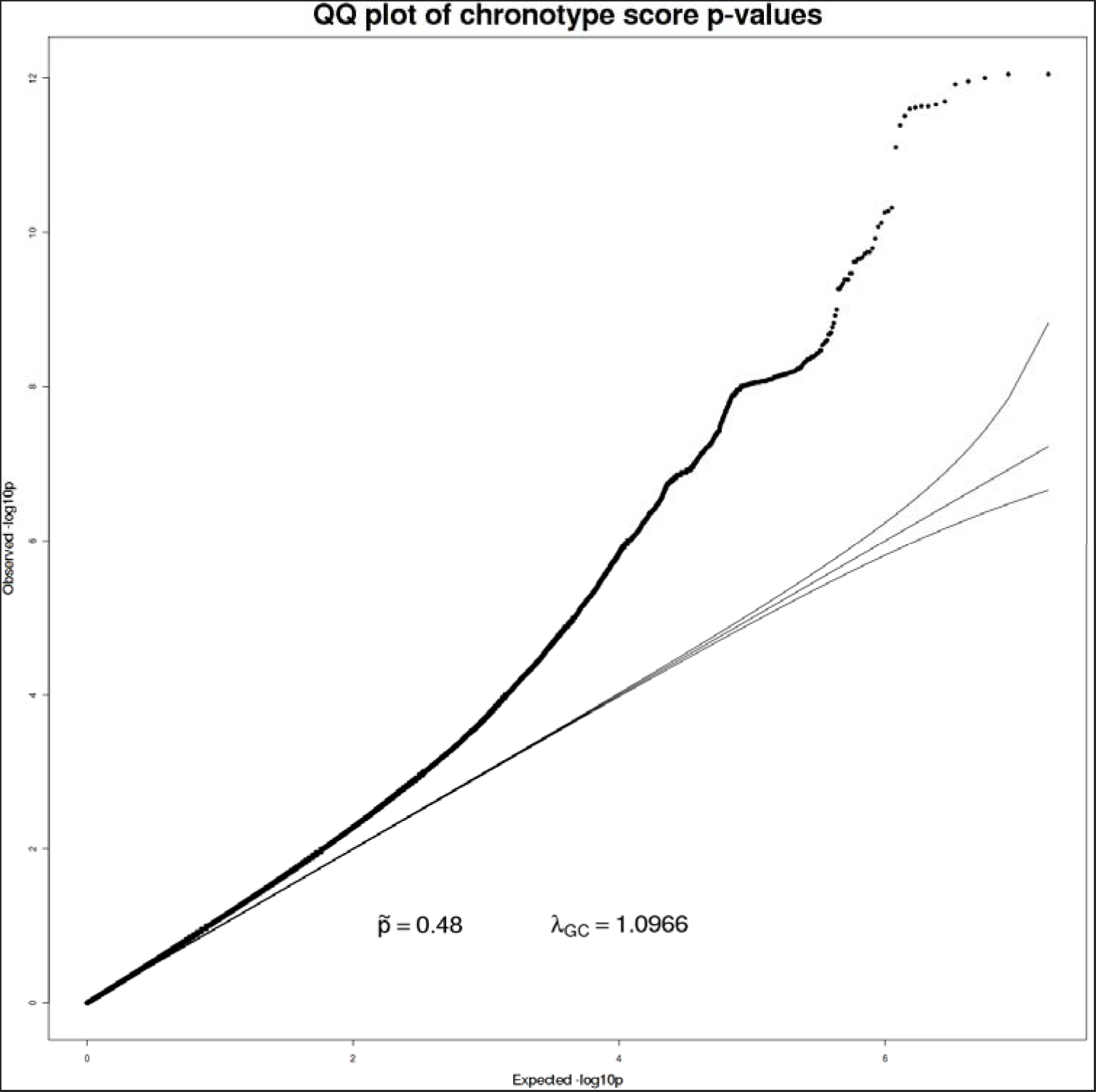
a) Manhattan and b) quantile-quantile (QQ) plot of chronotype score (inverse-normalised) P-values.

**Table 1.**
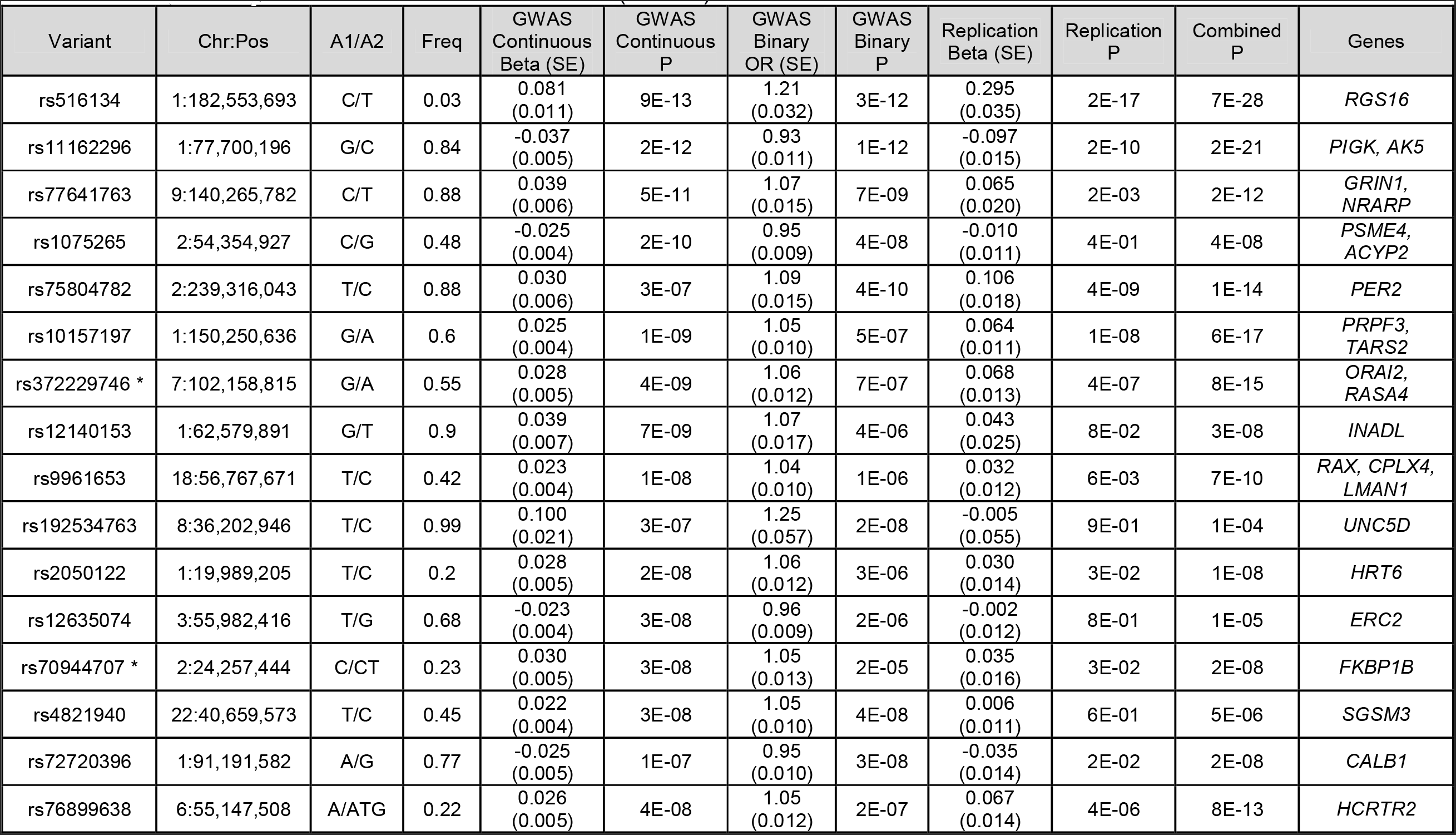
Genetic variants associated with chronotype. Sixteen loci associated with chronotype as either a continuous or binary trait in UK Biobank. Genes listed are candidate or nearest genes within 250Kb of the lead SNP. Odds ratios correspond to risk of morningness over eveningness. Beta, OR and frequency refers to A1. Replication data is based on continuous data and because the replication beta is in different units to the discovery GWAS beta a P-value meta-analysis was performed. * Proxies used for replication cohort: rs4729854 for rs372229746 (r^2^=0.33), and rs12621152 for rs70944707 (r^2^=0.33).

### Replication and validation of chronotype associations

Analysing UK Biobank data with that from 23andMe provides strong evidence that at least 13 of the 16 are robustly associated with chronotype. Thirteen of the chronotype signals remained at *P*<5x10^-8^ in a meta-analysis including UK Biobank and 89,283 individuals from 23andMe (Hu *et al*. Nature Communications, *In Press*), of which eleven reached *P*<0.05 in the same direction in 23andMe alone, and 15 of the 16 UK Biobank signals were in the same direction (binomial *P*=0.0002) (**Table 1**). We also attempted to validate the associations in 6,191 European-Ancestry from the Chronogen consortium and 2,532 Korean Ancestry individuals from the Insomnia, Chronotype and sleep EEG (ICE) consortium that used “Gold standard” chronotype questionnaire (Munich Chronotype Questionnaire – MCTQ and Morningness-Eveningness Questionnaire - MEQ). Given the sample size of 5% of the discovery UK Biobank study we assessed directional consistency rather than testing for replication P-values <0.05 or 0.05/16. In the European-Ancestry individuals 11 of the 16 signals were represented. Nine of these 11 variants had the same direction of effect as the discovery UK Biobank cohort (binomial test *P*=0.03) and one replicated at Bonferroni significance (rs12140153, *P*=0.003). In the Korean study, 9 signals were represented, four of which had the same direction of effect as the discovery UK Biobank cohort (binomial test *P*=1.00). The level of directional consistency in these two smaller studies is consistent with what would be expected in cohorts <5% the size of our discovery cohort.

### The chronotype-associated variants occur near genes known to be important in photoreception and circadian rhythms

The variant most strongly associated with chronotype, rs516134 (OR for morningness=1.21, [95% CI: 1.16,1.26], binary *P*=3.7x10^-12^, continuous *P*=8.9x10^-13^) occurs near *RGS16*, which is a regulator of G-protein signalling and has a known role in circadian rhythms ^14^ (**Table 1** **and** **Figure 2**). Another signal occurs near *PER2* (lead variant rs75804782, odds ratio=1.09, [95% CI: 1.06, 1.12], binary *P*=7.2x10^-10^, continuous *P*=3.2x10^-7^; **Figure 3**). *PER2* is a well-known regulator of circadian rhythms ^15-20^ and contains a variant, rs75804782, recently shown to be associated with iris formation ^21^ that is in LD (r^2^ = 0.65, D’ = 0.97) with our reported lead SNP. We also identified an association with a missense variant (rs12140153, OR=1.07 (95% CI: 1.04, 1.11), binary *P*=5x10^-6^, continuous *P*=7x10^-9^) in *INADL* (*InaD-like*) that encodes a protein thought to be important in organising and maintaining the “intrinsically photosensitive retinal ganglion cells”, cells that are known to communicate directly with the suprachiasmatic nucleus; the primary circadian pacemaker in mammals ^22^. As there is a reported link between season and reported chronotype ^23^, we carried out a sensitivity analysis in which we adjusted for month of attendance (to assessment centre); all associations remained genome-wide significant for the reported variants. We tested for enrichment of specific biological and molecular pathways using MAGENTA ^24^ but none had a clear link to circadian rhythms (**Supplementary Table 1**).

**Figure 2:**
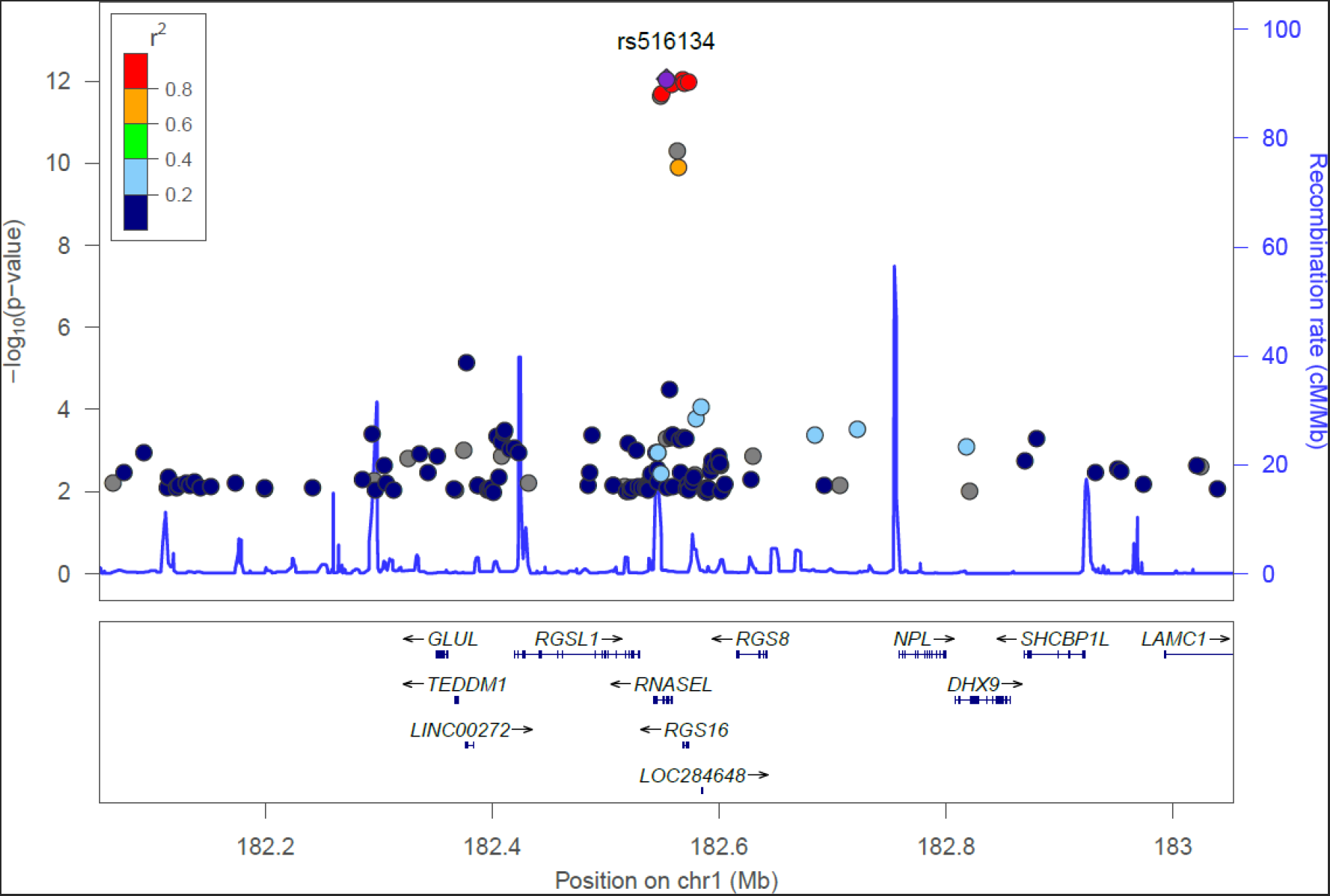
Locuszoom plot around *RGS16* locus.

**Figure 3:**
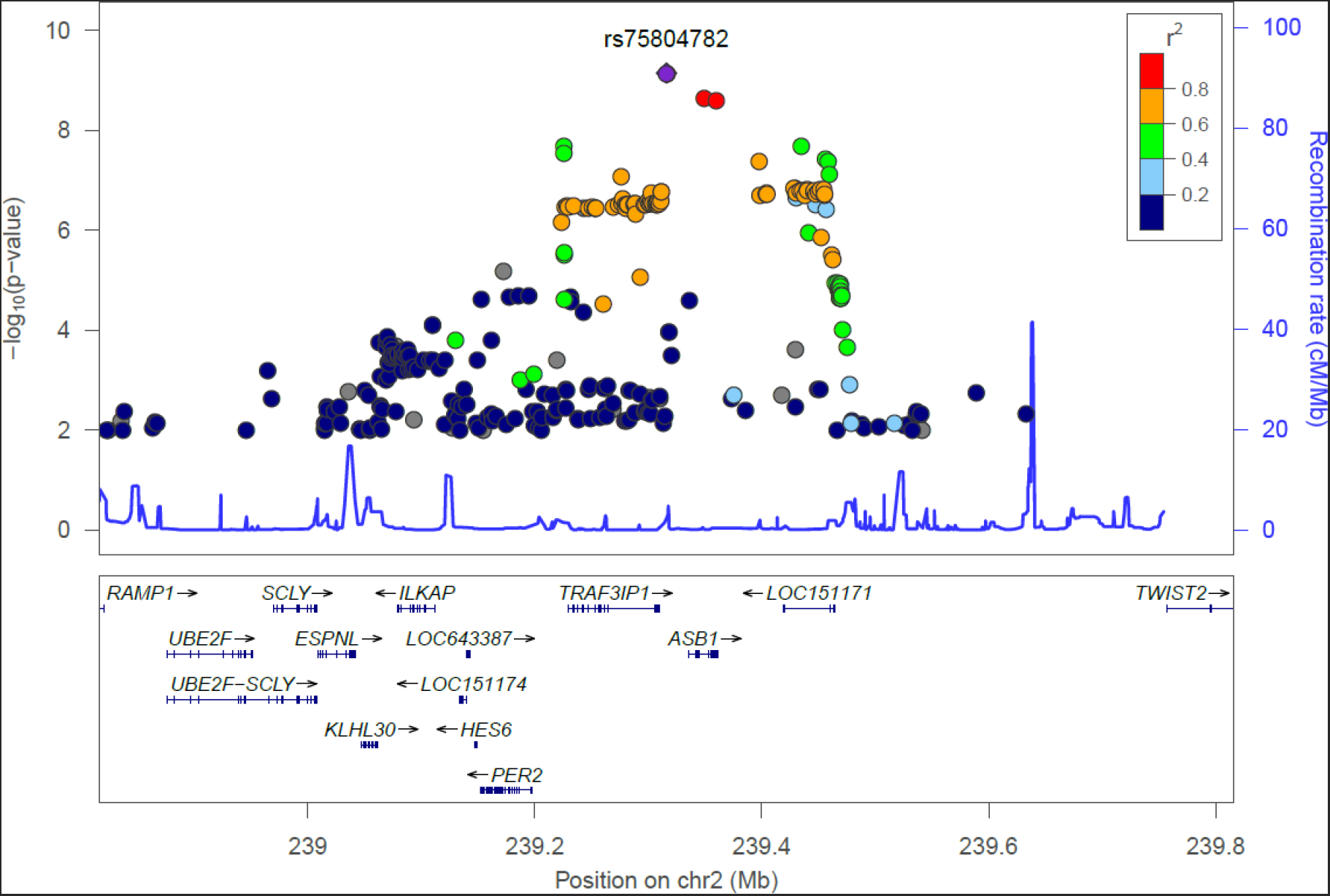
Locuszoom plot around *PER2*.

### Three loci associated with sleep duration

We performed genome-wide association studies on a binary sleep phenotype and a continuous sleep duration score for 16,761,225 imputed variants. **Figure 4** presents the overall results for these GWAS. Three loci reached genome-wide significance. The most strongly associated variant was rs62158211 with an average 2.6 minute (95% CI: 1.9 to 3.2 minutes, *P*=5.7x10^-16^) per-allele change in sleep duration and occurs at the previously reported association signal near *PAX8* ^13^. We identified two, novel, conditionally independent, signals that were located ∼900kb apart, one upstream and the other downstream of *VRK2*. The downstream variant, rs17190618, has an average per allele effect of 2.0 minutes (95% CI: 1.3 to 2.7 minutes), *P*=1.2x10^-9^, on sleep duration. The upstream variant, rs1380703 (which is not correlated with rs17190618, r^2^=0.002), has an average per allele effect of 1.6 minutes (95% CI: 1.1 to 2.2 minutes), *P*=7.6x10^-9^, on sleep duration. On adjusting for month of assessment, we saw marginally stronger associations for both rs62158211 (*P*=3x10^-16^) and rs1380703 (*P*=6x10^-9^), with no change for rs17190618. **Table 2** shows the three sleep duration loci and their lead variants. **Figure 5** shows locus zoom plots of the *VRK2* association signals. We did not replicate the association of a previously reported variant in *ABCC9* ^12^ with sleep duration (rs11046205, 0.1mins [95% CI: -0.6 to 0.7 minutes], *P*=0.83).

**Figure 4:**
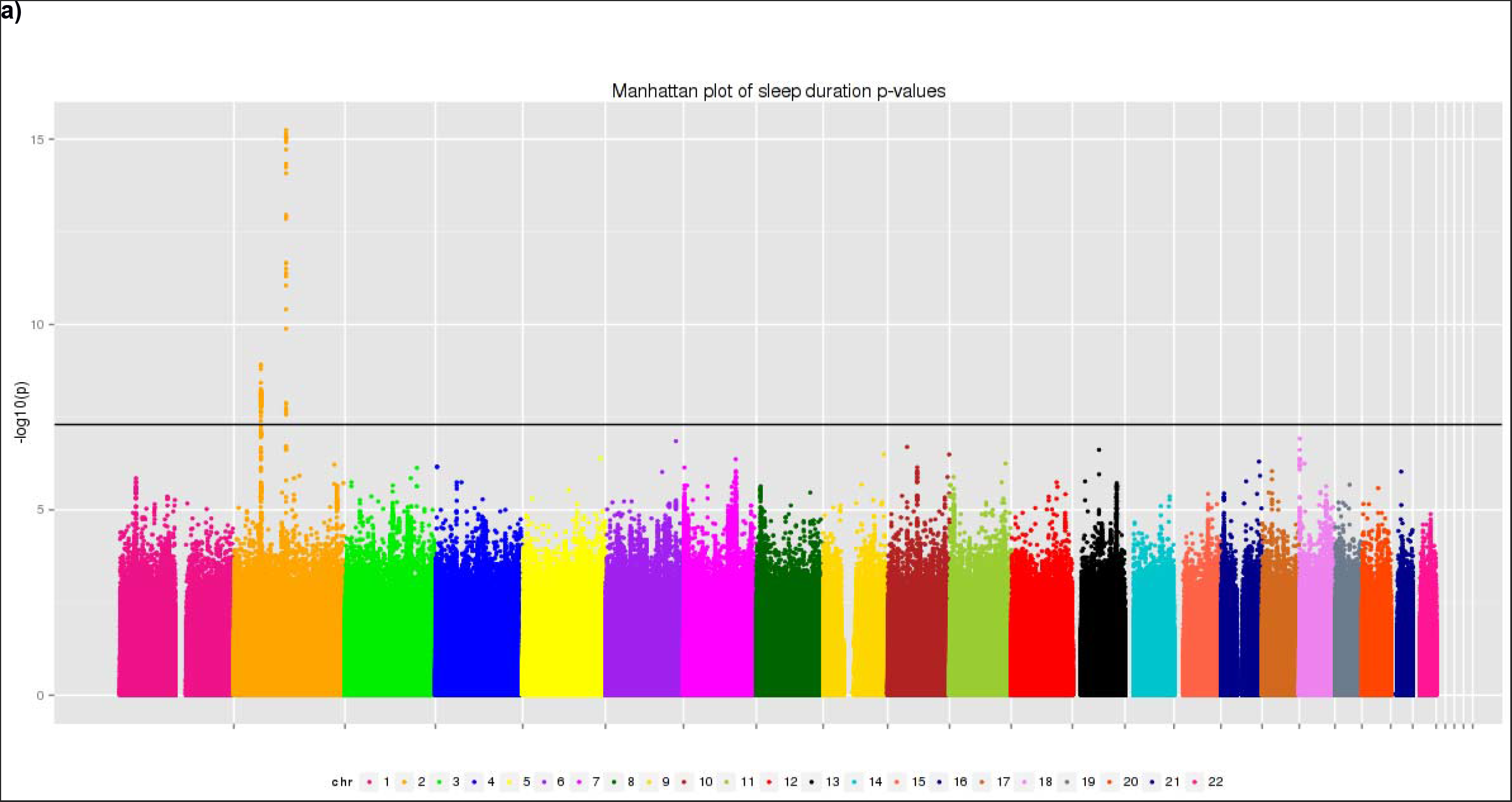
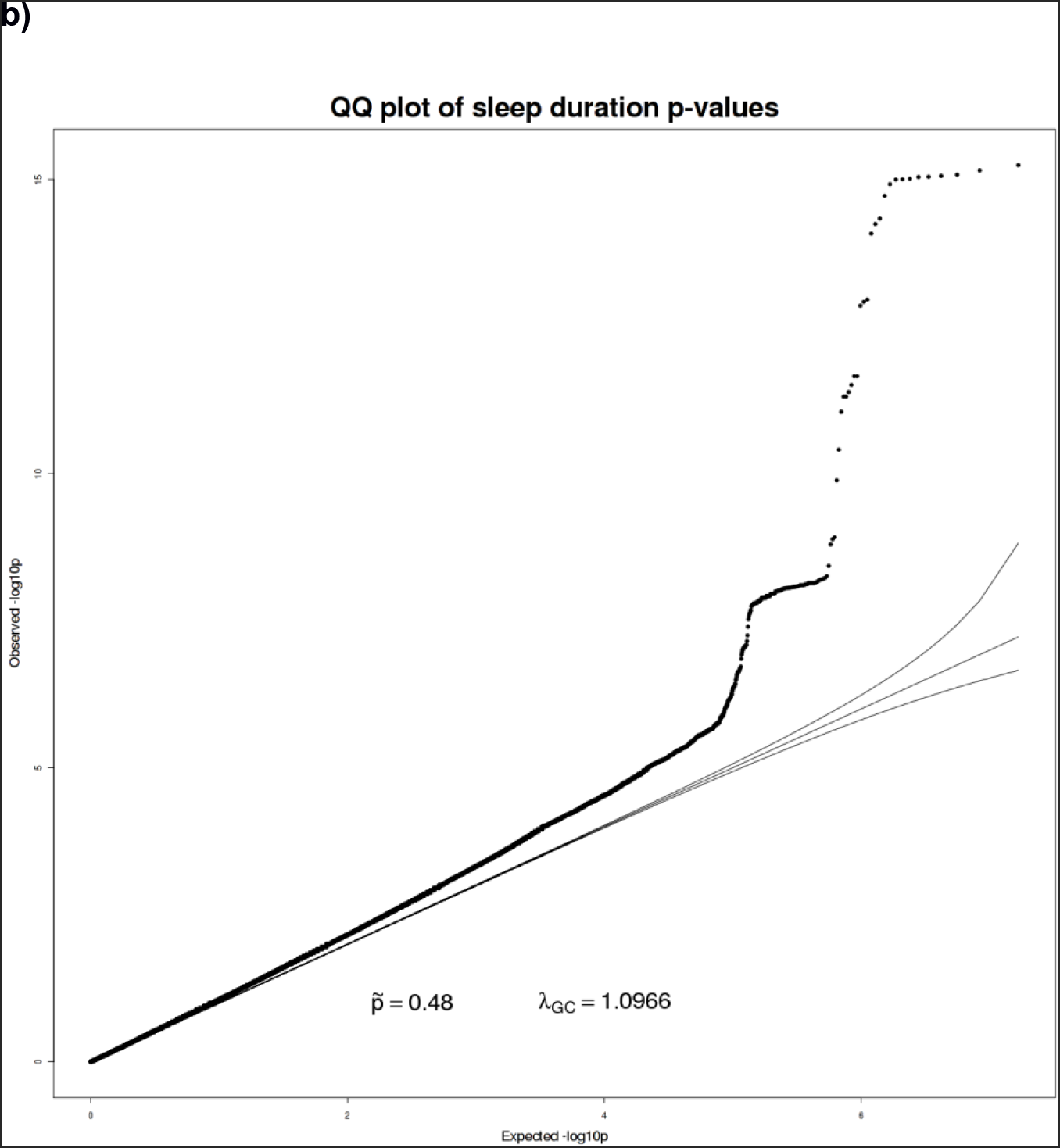
a) Manhattan and b) quantile-quantile plot of hours slept (inverse-normalised) P-values.

**Figure 5:**
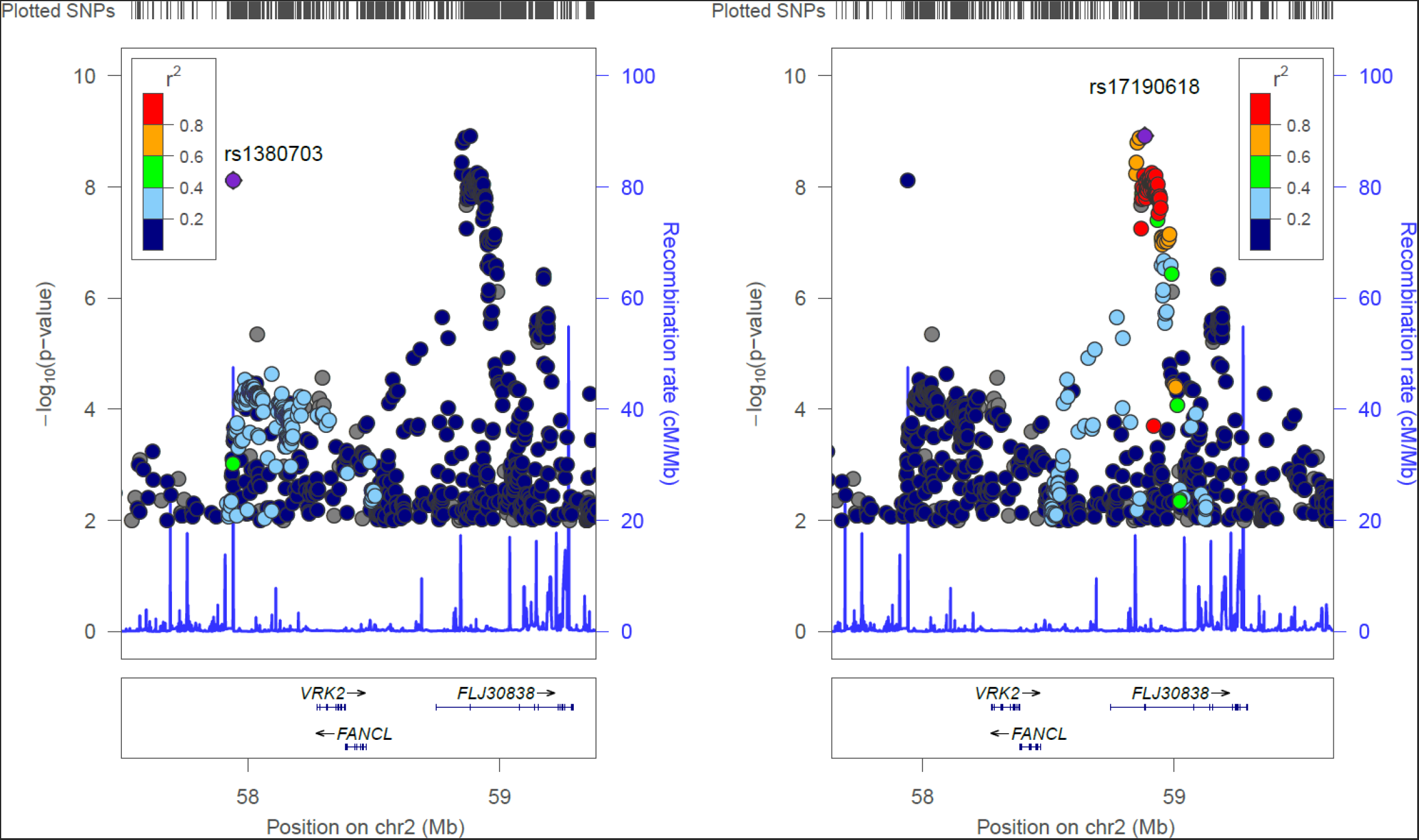
Locuszoom plots for the novel sleep duration associations at the *VRK2* locus.

**Table 2.** Three loci associated with sleep duration and their lead variants. Genes listed are candidate genes at each locus. Beta, OR and frequency refers to A1. Because the replication beta is in different units to the discovery GWAS beta a P-value meta-analysis was performed. Beta units are in hrs.

| Variant | Chr:Pos | A1/A2 | A1 Freq | GWAS<br>Continuous<br>Beta (SE) | GWAS<br>Continuous<br>P | GWAS<br>Binary<br>OR (SE) | GWAS<br>Binary P | Replication<br>Beta (SE) | Replication<br>P | Combined<br>P | Gene |
| --- | --- | --- | --- | --- | --- | --- | --- | --- | --- | --- | --- |
| rs62158211 | 2:114,106,139 | G/T | 0.79 | -0.039<br>(0.005) | 6E-16 | 0.94<br>(0.011) | 1E-07 | -0.053<br>(0.009) | 4E-9 | 2E-23 | PAX8 |
| rs17190618 | 2:58,882,765 | A/T | 0.84 | -0.033<br>(0.005) | 1E-09 | 0.96<br>(0.013) | 3E-04 | -0.035<br>(0.011) | 1E-3 | 5E-12 | VRK2 |
| rs1380703 | 2:57,941,287 | A/G | 0.62 | 0.025<br>(0.004) | 8E-09 | 1.06<br>(0.011) | 8E-08 | 0.021<br>(0.008) | 1E-2 | 3E-10 | VRK2 |

### Replication of novel sleep duration hits

To replicate the two novel sleep duration hits we used data from 47,180 individuals from a published study ^13^. The variant rs17190618 replicated with effect size=2.1 minutes (95% CI:0.8 to 3.3), *P*=0.001, meta-analysis *P*=5x10^-12^. The variant rs1380703 replicated with effect size=1.3 minutes (95% CI: 0.3 to 2.2), *P*=0.01, meta-analysis *P*=3x10^-10^).

### Sleep duration and chronotype are heritable and genetically correlated with BMI, Type 2 diabetes and psychiatric disease

Using LD-score regression we estimated the heritability of chronotype and sleep duration within UK Biobank to be 0.12 (0.007), and 0.07 (0.007), respectively. There was no significant genetic correlation between sleep duration and chronotype (rG=0.0177, *P*=0.70). Chronotype was nominally genetically correlated with BMI (rG=0.056, *P*=0.048), but not Type 2 diabetes (rG=0.004, *P*=0.99). As the relationship between sleep duration with BMI and risk of T2D is U-shaped (see **Supplementary Figure 1**), we defined two further binary phenotypes; undersleepers (<7 vs. 7-8 hours) and oversleepers (>8 vs. 7-8 hours). There was a strong genetic correlation between undersleeping and BMI (rG=0.147, *P*=1x10^-5^), but not T2D (rG=0.022,*P*=0.79). There was also a genetic correlation between oversleeping and both BMI (rG=0.097, *P*=0.039) and T2D (rG=0.336, *P*=0.001). We also performed LD-score regression analyses against a range of other diseases and traits where GWAS summary statistics are publically available (**Supplementary Table 2**). Schizophrenia was genetically correlated (after adjusting for the number of tests) with hours slept (rG=0.25, *P*=1x10^-4^), oversleeping (rG=0.32, *P*=9x10^-4^), but not significantly correlated with undersleeping (rG=-0.11, *P*=0.097).

### Mendelian randomisation analyses provide no consistent evidence that higher BMI affects self-reported morningness or vice-versa

Using a genetic risk score of 69 known BMI variants ^25^ (listed in **Supplementary Table 3**) as an instrumental variable, we next performed Mendelian randomisation analyses in the UK Biobank study to test the potential causal role of BMI in chronotype and sleep. Instrumental variables analyses using variants and their effect sizes identified by previous studies ^25^ provided no consistent evidence that self-reported “morningness” causally affects BMI or risk of type 2 diabetes (**Supplementary Table 4**). Association statistics of the BMI variants with chronotype are given in **Supplementary Table 3**. We repeated these analyses using a genetic risk score consisting of 55 type 2 diabetes SNPs ^26^ and did not find any evidence of causality. Association of the chronotype-associated variants with BMI are given in **Supplementary Table 5**. Performing the reciprocal Mendelian randomization analysis using a genetic risk score of the 13 replicated chronotype variants, with effect sizes obtained from 23andMe, we found no consistent evidence in the UK Biobank data that morningness or eveningness leads to higher BMI (**Supplementary Table 4**).

### No evidence that BMI and Type 2 diabetes are causally associated with sleep duration

Using the same genetic risk score of 69 known BMI variants as an instrument, we saw no consistent evidence that higher BMI increased an individual’s likelihood of being an undersleeper (IVreg2 *P*=0.95, IVW *P*=0.05) or an oversleeper (IVreg2 *P*=0.29, IVW *P*=0.62) in the UK Biobank data (**Supplementary Table 4**). Because there were only three genetic variants of small effect associated with sleep duration, we did not perform any Mendelian Randomisation analyses of sleep on BMI or type 2 diabetes risk.

## Discussion

We performed a genome-wide association study of sleep duration and morningness in 128,266 individuals from the UK Biobank study. We discovered and replicated two novel loci associated with sleep duration. Through replication in a study of 89,823 individuals from 23andMe we found 13 genome-wide significant loci for chronotype. These loci occur in or near circadian rhythm and photoreception genes and provide new insights into circadian rhythm and sleep biology and their links to disease.

The two novel sleep duration association signals that we have discovered and replicated in this study occur upstream and downstream of *VRK2* (*vaccinia related kinase 2*). VRK2 is a serine/threonine kinase important in several signal transduction cascades, and variants near *VRK2* are associated with schizophrenia ^27^ and epilepsy ^28^. The two sleep duration variants we identified do not represent the same signals as those associated with schizophrenia at genome wide significance but one is associated with schizophrenia (based on publically available data from the schizophrenia genetics consortium (rs1380703 *P*=2x10^-5^), with the allele associated with more sleep being associated with higher risk of schizophrenia). Furthermore, the variants associated with epilepsy and schizophrenia at genome wide significance are associated with sleep duration in UK Biobank (epilepsy lead variant rs2947349 ^28^, *P*=2x10^-5^ and schizophrenia lead variant ^27^ rs11682175 *P*=3x10^-5^) but did not reach genome wide significance. We also observed genetic correlation between sleep duration and schizophrenia using LD-score regression (rG=0.25, *P*=1x10^-4^). Further work is required to determine whether variation in *VRK2* either has independent associations with both sleep and schizophrenia or whether there is some causal link between sleep duration and schizophrenia and epilepsy.

Several of the loci that we identified as associated with chronotype contain genes that have a known role in circadian rhythms. The most strongly associated variant, rs516134, occurs 20kb downstream of *RGS16* (regulator of G protein signalling 16). *RGS16* has recently been shown to have a key role in defining 24 hour rhythms in behaviour ^14^. In mice, gene ablation of *Rgs16* lengthens the circadian period of behavioural rhythm ^14^. By temporally regulating cAMP signalling, Rgs16 has been shown to be a key factor in synchronising intercellular communication between pacemaker neurons in the suprachiasmatic nucleus (SCN), the centre for circadian rhythm control in humans.

The association signal with lead SNP rs75804782 occurs ∼100kb upstream of *PER2* (*Period 2*). Per2 is a key regulator of circadian rhythms and is considered one of the most important clock genes, and, under constant darkness, *Per2* knockout mice show arrhythmic locomotor activity ^15-20^. This locus also contains a variant that has recently been shown to be associated with iris furrow contractions ^21^. Our signal is very likely to represent the same association and suggests a link between iris function and chronotype (rs75804782 has an LD r^2^ = 0.65 and D’ = 0.97 with the reported lead SNP, rs3739070). Larsson *et al*. ^21^ suggest *TRAF3IP1* as the most likely candidate gene at the locus because of its critical role in the cytoskeleton and neurogenesis. Further work is needed to elucidate whether the chronotype association at this locus acts through *PER2* or *TRAF3IP1*.

Several of the variants associated with chronotype are also associated with BMI and we found genetic correlation between chronotype and sleep duration and BMI. There is substantial evidence for a role of sleep disruption and circadian rhythms in metabolic disease ^1^. Data from animal models and epidemiology provide strong evidence that sleep quality or disrupted circadian rhythms can cause metabolic diseases including obesity and type 2 diabetes ^4-6, 8-10^. Our Mendelian Randomisation analyses provided no consistent evidence for a role of higher BMI leading to increased self-reported morningness.

There are some important limitations to our study. First, chronotype and sleep duration were self-reported and are subject to reporting bias (e.g. obese individuals may be more likely to falsely claim to be morning people). Second, whilst we did not find any evidence that overall chronotype or sleep duration causally lead to obesity or type 2 diabetes, it is possible that sub-pathways of genes involved in, for example, feeding behaviour may be important in both obesity and chronotype regulation. The availability of the full UK Biobank study of 500,000 will provide further insight into this relationship.

In conclusion, we have identified novel genetic associations for chronotype and sleep duration. The chronotype loci cluster near genes known to be important in determining circadian rhythms and will provide new insights into circadian regulation. Our results provide new insights into circadian rhythm and sleep biology and their links to disease.

## Materials and Methods

### Discovery Samples

We used 128,266 individuals of British descent from the first UK Biobank genetic data release (see http://biobank.ctsu.ox.ac.uk). British-descent was defined as individuals who both self-identified as white British and were confirmed as ancestrally Caucasian using principal components analyses (http://biobank.ctsu.ox.ac.uk). Of these individuals, 120,286 were classified as unrelated, with a further 7,980 first- to third-degree relatives of these. As the association tests were carried out in BOLT-LMM ^29^, which adjusts for relationships between individuals and corrects for population structure, we included all 128,266 related white British individuals in the association analyses.

### Genotyping and quality control

We used imputed variants provided by the UK Biobank. Details of the imputation process are provided at the UK Biobank website (see http://biobank.ctsu.ox.ac.uk). For this study we only included the ∼16.7M imputed variants with an imputation R^2^ ≥ 0.4, MAF ≥ 0.001 and with a Hardy–Weinberg equilibrium *P*>1x10^-5^.

### Phenotypes

#### Chronotype

UK Biobank provides a single measure of Chronotype, from which we produced a continuous and a dichotomous phenotype. Chronotype (or morningness) is a self-reported measure and asks individuals to categorise themselves as “Definitely a ‘morning’ person”, “More a ‘morning’ than ‘evening’ person”, “More an ‘evening’ than a ‘morning’ person”, “Definitely an ‘evening’ person” or “Do not know”, which we coded as 2, 1, -1, -2 and 0 respectively, in our raw continuous “score”. Individuals had the option not to answer; these individuals were set to missing. We then produced a normally distributed phenotype by adjusting the raw phenotype for age, gender and study centre (categorical) and inverse normalising the resulting residuals. The dichotomous chronotype trait defines morning people (“Definitely a ‘morning’ person” and “More a ‘morning’ than ‘evening’ person”) as cases and evening people (“Definitely an ‘evening’ person” and “More an ‘evening’ than a ‘morning’ person”) as controls. All other individuals are coded as missing. All results reported for continuous chronotype refer to the inverse-normalised residualised chronotype “score”. For interpretable results, however, we report effect sizes using the odds ratios of the dichotomous chronotype phenotype. A total number of 127,898 and 114,765 individuals were available with non-missing continuous and binary chronotype phenotypes, respectively, for the association tests; for the Mendelian Randomisation this became 119,935 and 107,634 respectively.

#### Sleep duration

The UK Biobank also provides self-reported “sleep duration”, in which individuals were asked to provide the average number of hours slept in a 24-hour period. The phenotype was derived by first excluding individuals reporting greater than 18 hours sleep, then adjusting for age, gender and study centre (categorical) and obtaining the model residuals and finally inverse-normalising to assure a normally distributed phenotype. When reporting results for the continuous sleep duration phenotype, we are referring to the inverse-normalised phenotype, though we report effect sizes of the residualised phenotype to allow easier interpretation of results. There were 127,573 individuals with reported sleep duration available for the association tests, with 119,647 available for the MR analyses.

#### “Oversleepers” and “Undersleepers”

These two dichotomous phenotypes share the same set of controls; those individuals that reported sleeping either 7 or 8 hours (81,204 individuals). In oversleepers, cases (10,102 individuals) are those reporting 9 or more hours sleep on average, whereas undersleeper cases (28,980 individuals) are those reporting 6 or fewer hours.

#### BMI

The UK Biobank provided a BMI (weight (kg)/height^2^) measurement and an estimate based on electrical impedance analyses. To help avoid reporting error we excluded individuals with significant differences (>4.56 SDs) between these two variables where both were available. If only one of these measurements was available this was used. We corrected BMI by regressing age, sex, study centre, and the first 5 within-British principal components and taking residual values. We then inverse normalised the residuals. A total of 119,684 white-British individuals with BMI and genetic data were available for the Mendelian Randomisation analyses.

#### Type 2 diabetes

Individuals were defined as having T2D if they reported either T2D or generic diabetes at the interview stage of the UK Biobank study. Individuals were excluded if they reported insulin use within the first year of diagnosis. Individuals reportedly diagnosed under the age of 35 years or with no known age of diagnosis were excluded, to limit the numbers of individuals with slow-progressing autoimmune diabetes or monogenic forms. Individuals diagnosed with diabetes within the last year of this study were also excluded as we were unable to determine whether they were using insulin within this time frame. A total of 4,040 cases and 113,735 controls within the white British subset of UK Biobank were identified with genetic data available.

### Genome-wide association analysis

To perform the association tests, we used BOLT-LMM ^29^ to perform linear mixed models (LMMs) in the 128,266 individuals. We used BOLT-LMM as it adjusts for population structure and relatedness between individuals whilst performing the association tests with feasible computing resources. As it adjusts for population structure and relatedness between individuals whilst performing the association tests, it allowed us to include the additional 7,980 related individuals and therefore improved our power to detect associations. To calculate the relationships between individuals, we provided BOLT-LMM a list of 328,928 genotyped SNPs (MAF>5%; HWE *P*>1x10^-6^; missingness<0.015) for the individuals included in the association analysis and used the 1000 Genomes LD-Score table provided with the software.

As the continuous phenotypes were derived by adjusting for age, gender and study centre, the LMM only included chip (BiLEVE vs. UKBiobank arrays) as a covariate at run-time (see http://www.ukbiobank.ac.uk/wp-content/uploads/2014/04/UKBiobank_genotyping_QC_documentation-web.pdf). The binary phenotypes were unadjusted and so included age, gender and chip at run-time. BOLT-LMM reported no improvement of the non-infinitesimal mixed model test over the standard infinitesimal test and so all association results reported in this paper are for the infinitesimal model ^29^.

### Chronotype replication samples

Participants were from the customer base of 23andMe, Inc. The descriptions of the samples, genotyping and imputation are in Hu *et al*. Nature Communications, *In Press*. Of the 16 chronotype-associated variants for which we attempted replication, 10 were available from imputation from the 1000 Genomes imputation panel phase 1 pilot. An additional 4 were imputed from the phase 1 version 3 1000 Genomes imputation panel. The final two could not be imputed. We used http://analysistools.nci.nih.gov/LDlink/ to find proxies --the best available were rs4729854 for rs372229746 (r2=0.33), and rs12621152 for rs70944707 (r2=0.33). We meta-analysed P-values from the discovery and replication samples using sample size weighting implemented in METAL ^30^.

### Chronotype validation samples

Genotypes consisting of both directly typed and imputed SNPs were used for the individual GWAS ^12^. To avoid over-inflation of test statistics due to population structure or relatedness, we applied genomic control for the independent studies and meta-analysis. Linear regression for associations with normalized chronotype was performed (see ^12^ for packages used) under an additive model, with SNP allele dosage as predictor and with age, age^2^, gender, normalized sleep duration, season of assessment (dichotomized based on time of the year, and day-light savings time – DST or standard zone time assessments) as covariates. A fixed-effects meta-analysis was conducted with GWAMA using the inverse-variance-weighted method and low imputation quality (Rsq/proper_info < 0.3) were dropped from the meta-analysis.

### Pathway analyses

Pathway analyses were carried out in MAGENTA using all available libraries provided with the software. We included all imputed variants with association *P*<1x10^-5^ from the continuous chronotype trait. For the results presented in **Supplementary Table 1**, we used gene upstream and downstream limits of 250Kb, excluded the HLA region (default setting) and set the number of permutations for GSEA estimation at 10,000 (default).

### Genetic correlation analyses

Genetic correlations (see ^31^ for methodology) between traits were calculated using the LD Score Regression software LDSC (available at https://github.com/bulik/ldsc/) ^32^. Summary statistics of our traits outputted by BOLT-LMM were first “munged”, a process that converts the summary statistics to a format that LDSC understands and aligns the alleles to the Hapmap 3 reference panel, removing structural variants and multi-allelic and strand-ambiguous SNPs. Genetic correlations were then calculated between our phenotypes and a set of 100 phenotypes for which summary statistics are publicly available (full list in **Supplementary Table 2**). We used precomputed LD structure data files specific to Europeans of HAPMAP 3 reference panel, obtained from (http://www.broadinstitute.org/∼bulik/eur_ldscores/) as suggested on the LDSC software page.

### Mendelian Randomisation IV analysis

The 13 variants in **Table 1** which reached *P*<5x10^-8^ in combined analyses were used as chronotype instruments in the Mendelian Randomisation analyses. Where binary and continuous traits shared a locus, we selected the top variant of the continuous trait over that of the binary. For loci that reach GW-significance in the binary trait only, we selected the top variant but used the effect size from the continuous trait.

To test for a causal effect of BMI on chronotype and sleep-duration, we selected 69 of 76 common genetic variants that were associated with BMI at genome wide significance in the GIANT consortium in studies of up to 339,224 individuals (**Supplementary Table 3**) ^25^. We limited the BMI SNPs to those that were associated with BMI in the analysis of all European ancestry individuals and did not include those that only reached genome-wide levels of statistical confidence in one-sex only, or one stratum only. Variants were also excluded if known to be classified as a secondary signal within a locus. Three variants were excluded from the score due to potential pleiotropy (rs11030104 [*BDNF* reward phenotypes], rs13107325 [*SLC39A8* lipids, blood pressure], rs3888190 [*SH2B1* multiple traits]), three due to being out of HWE (rs17001654, rs2075650 and rs9925964) and the last variant due to not being present in the imputed data (rs2033529).

For testing reverse causality of type 2 diabetes on our sleep phenotypes, we used 55 of 65 common variants (listed in **Supplementary Table 3**) known to be associated with type 2 diabetes at genome wide significance in a meta-analysis of 34,840 cases and 114,981 ^26^, excluding those known or suspected to be pleiotropic.

We performed the Mendelian Randomisation analysis two ways; firstly using instrumental variables (IV) using STATA’s “IVreg2” function ^33^ and secondly through the inverse-variance weighted (IVW) and MR-Egger methods described in ^34^. Analyses were performed in STATA 13.1 (StataCorp. 2013. Stata Statistical Software: Release 13. College Station, TX: StataCorp LP.).

In the instrumental variables method, we generated genetic risk scores (GRS) for BMI and type 2 diabetes using the published list of associated variants and their respective betas. For Chronotype, we generated a GRS using the thirteen replicated variants and their respective betas from 23andMe summary statistics. Using the IVreg2 command, we performed two-stage least squares estimation to calculate the effect of predicted exposure (through the GRS) on the continuous outcome traits. For binary outcomes (type 2 diabetes, undersleeper and oversleeper), we manually carried out the two-stage process by regressing the exposure trait on its GRS and storing both predicted values and residuals. We then used these predicted values and residuals as independent variables in a logistic regression where the dependent variable was the binary outcome.

The inverse-variance weighted (IVW) method is equivalent to a meta-analysis of the associations of the individual instruments and uses associations between the instruments and both the exposure and the outcome to estimate the additive effect of the instruments combined ^34^. The MR-Egger method is a modification to the IVW method that allows the inclusion of “invalid” instruments (i.e. those that don’t satisfy all three conditions), by performing Egger regression using the summary data of the variants. The IVW and Egger methods operate under the assumption that all instruments are valid, in that they satisfy the three IV conditions: the genetic variants are 1) independent of confounders, 2) associated with the exposure and 3) independent of the outcome. The MR-Egger method, however, accounts for the fact that genetic variants could be pleiotropic and may influence the outcome via pathways other than through the exposure and therefore the resulting association between genetic instruments and the outcome should not be biased by invalid instruments and pleiotropy. The MR-Egger method was used purely as a sensitivity test for the IVW method and so MR-Egger results were not considered if the IVW result did not reach nominal significance.

For the IVW and MR-Egger methods, associations of genetic instruments (variants) with both exposure and outcome phenotypes were generated in STATA by regressing the phenotype against the instrument while adjusting for covariates. As a further sensitivity test, we also repeated these analyses by replacing exposure phenotype-variant associations with their respective published betas and found only slight differences in betas and P-values, though all exposure-outcome associations remained non-significant.

## Acknowledgements

This research has been conducted using the UK Biobank Resource. We would like to thank the research participants and employees of 23andMe for making this work possible.

**Funding Information**. S.E.J. is funded by the Medical Research Council (grant: MR/M005070/1) M.A.T., M.N.W. and A.M. are supported by the Wellcome Trust Institutional Strategic Support Award (WT097835MF). A.R.W., T.M.F and H.Y. are supported by the European Research Council grants: SZ-245 50371-GLUCOSEGENES-FP7-IDEAS-ERC and 323195. R.M.F. is a Sir Henry Dale Fellow (Wellcome Trust and Royal Society grant: 104150/Z/14/Z). R.B. is funded by the Wellcome Trust and Royal Society grant: 104150/Z/14/Z. J.T. is funded by a Diabetes Research and Wellness Foundation Fellowship. This study was provided with biospecimens and data from the Korean Genome Analysis Project (4845-301), the Korean Genome and Epidemiology Study (2010-E71001-00, 2011-E71004-00, and 2011-E71008-00), and Korea Biobank Project (4851-307) that were supported by the Korea Centers for Disease Control & Prevention, Republic of Korea.The funders had no influence on study design, data collection and analysis, decision to publish, or preparation of the manuscript.

**Duality of Interest**. No potential conflicts of interest relevant to this article were reported.

---

**Supplementary Figure S1:**
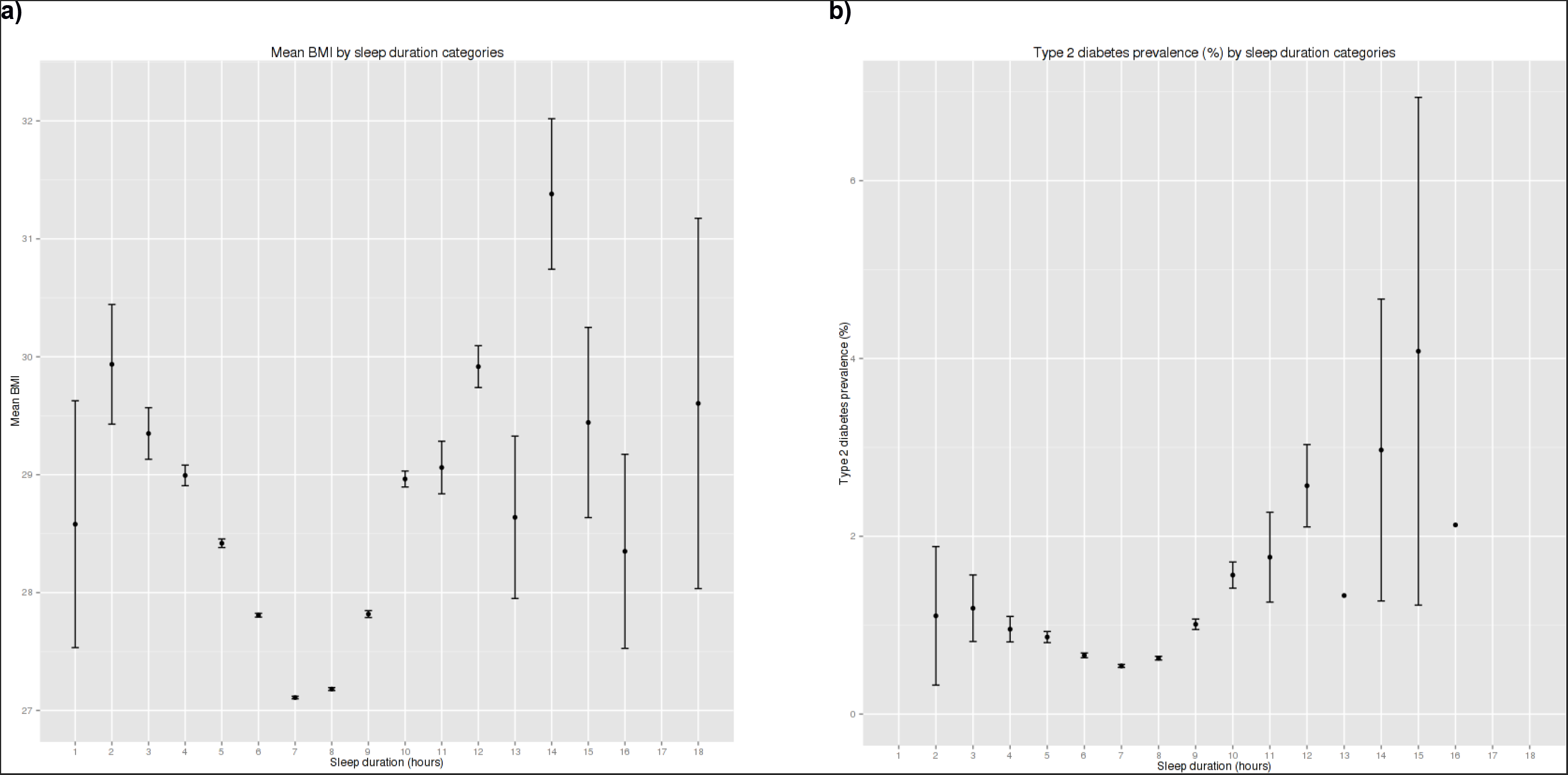
Distribution of a) self-reported BMI and b) proportion of individuals reporting type 2 diabetes in each of the possible sleep duration categories.

